# Dual single-nucleus gene expression atlas of grapevine and *Erysiphe necator* during early powdery mildew infection

**DOI:** 10.1101/2025.08.11.669584

**Authors:** Maria-Sole Bonarota, Jadran F. Garcia, Mélanie Massonnet, Mirella Zaccheo, Rosa Figueroa-Balderas, Noé Cochetel, Dario Cantu

## Abstract

We applied dual-organism single-nucleus transcriptomics to study the interaction between grapevine leaves and *Erysiphe necator,* the causal agent of powdery mildew, at one and five days post-infection, including controls and three biological replicates. We generated a grapevine leaf atlas encompassing over 100,000 nuclei, and a pathogen atlas of more than 3,000 nuclei. We successfully annotated all major grapevine cell types, including mesophyll, epidermis, phloem and xylem parenchyma, companion cells, and guard cells. We identified key *E. necator* structures, including appressoria, haustoria, and hyphae, and provided a list of novel cell type markers for both species. We reveal structure-specific gene expression programs in *E. necator* laying a foundation for future studies of fungal development and virulence mechanisms. In the host, we identified spatially distinct expression patterns of defense-related genes. As the infection progressed, we observed the activation of a coordinated immune response involving multiple cell types, mainly epidermal and mesophyll cells. High-dimensional weighted gene co-expression network analysis identified key hubs and networks associated with cell type-specific signaling and defense response. We describe a spatial separation of pattern- and effector-triggered immunity, supporting a model in which pattern-triggered immunity is activated at the site of pathogen contact and effector-triggered immunity is induced in surrounding tissue.

## Introduction

Grapevine powdery mildew (PM) is a devastating disease of grapes, particularly domesticated grapevines (*Vitis vinifera* L.). The disease is caused by the fungus *Erysiphe necator* Schw. [syn. *Uncinula necator* (Schw.) Burr.], a highly specialized biotrophic pathogen that colonizes the surface of above-ground green organs such as leaves, stems, flowers, and immature berries. Conidia dispersed by wind or humidity germinate on the grapevine cuticle, forming an appressorium that penetrates the epidermal wall and establishes a haustorium within the host cell (Qiu et al., 2015). Through this specialized organ *E. necator* acquires nutrients, while secreting proteins that manipulate host metabolism and suppress immune responses (Sree et al., 2024). The fungus spreads across the host surface through extensive hyphal growth, forming a visible mycelium (Gadoury et al., 2012; Qiu et al., 2015). Infected grapevine leaves exhibit reduced photosynthesis and hormonal imbalance (Kunova et al., 2021), while early berry infection can lead to berry cracking, with overall impacts on the crop including reduced yields, increased acidity, and lower sugar and anthocyanin content in mature fruit (Calonnec et al., 2004).

Grapevine defense against *E. necator* relies on penetration resistance and programmed cell death (PCD), which limit fungal access to nutrients and restrict colonization (Qiu et al., 2015). Pathogen-associated molecular pattern (PAMP)-triggered immunity (PTI) is the first layer of innate plant defense and is initiated when the host cell plasma membrane receptors (e.g., receptor-like kinase (RLK) proteins) recognize PAMPs, such as chitin, a major constituent of fungal cell wall (Boller and Felix, 2009). PTI activation leads to a mitogen-activated protein kinase (MAPK) cascade, triggering a broad range of defense responses, including transcriptional reprogramming, hormone signaling, oxidative burst, and cell wall strengthening (Meng and Zhang, 2013). Fungal pathogens can suppress PTI through the secretion of effector proteins, which can be detected by intracellular receptors (Tsuda and Katagiri, 2010), typically nucleotide-binding leucine-rich repeat proteins (NLR; Gururani et al., 2012). This recognition initiates the effector-triggered immunity (ETI), often associated with PCD.

Although much has been learned about *E. necator* biology and grapevine susceptibility through bulk transcriptomics and physiological studies (Fung et al., 2008; Amrine et al., 2015; Jiao et al., 2021; Sree et al., 2024), our understanding of the interaction remains limited by the lack of cellular resolution. Which grapevine cell types respond to infection, and how fungal structures coordinate their activity across tissues, remain open questions. Since epidermal cells are in direct contact with the pathogen, we hypothesize that they initiate the response to the pathogen (e.g., PTI and ETI), but whether, how, and when different cell types respond to the infection is unknown. The signaling processes that initiate metabolic changes to sustain the infected epidermis and the biotrophic fungus are likely to involve additional tissues, such as the mesophyll and the vascular tissue. Moreover, fungal gene expression dynamics across distinct infection structures, such as haustoria and appressoria remain unexplored. Answering these questions requires tools that can resolve host-pathogen interactions at cellular scale. Recent single-cell studies in model systems like *Arabidopsis thaliana* have shown that different host cell types mount distinct transcriptional responses during infection, even within the same organ (Delannoy et al., 2023; Nobori and Ecker, 2023; Tang et al., 2023; Zhu et al., 2023).

Here, we present the first application of single-cell transcriptomics to a grapevine–PM interaction. Using dual-organism single-nucleus RNA sequencing (snRNA-seq) to simultaneously profile host and pathogen gene expression, we investigate the transcriptional landscape of infected grapevine leaves at early stages of colonization by *E. necator*. Our study focuses on the interaction between a commercially important cultivar (Cabernet Sauvignon) and a highly virulent *E. necator* strain, with the goal of describing cell-type-specific host responses and fungal activity during colonization. In addition to revealing new aspects of grapevine responses to powdery mildew, our study lays the groundwork for future investigations of resistant genotypes, spatial regulation of immunity, and effector function. We also identify novel cell-type markers in both host and pathogen, providing tools for probing functional specialization and cellular reprogramming.

## Results

### Single-nucleus transcriptomics of powdery mildew-infected grapevine leaves

Grapevine leaves were inoculated with the virulent *E. necator* strain UCD-TS1. Pathogen development was monitored by scanning electron microscopy (SEM) at one and five dpi. At one dpi, appressoria and primary hyphae were observed on the leaf surface (**Fig. 1A**), indicative of the early stage of the biotrophic colonization. By five dpi, the pathogen had developed extensive mycelium with both primary and secondary hyphae, consistent with advanced stages of infection (**Fig. 1B)**. To support the selection of time points for single-nucleus RNA-seq, we first conducted bulk-tissue RNA-seq at one, five, and ten dpi to assess global transcriptional responses. We identified 2,269, 3,636, and 2,802 differentially expressed genes (DEGs) compared to the mock-inoculated controls at one-, five-, and ten-dpi, respectively (adjusted *p* value < 0.05), with 1,289, 2,181, and 1,424 upregulated (**Fig. S1A**) and 980, 1,455, and 1,378 downregulated (**Fig. S1B**). Venn diagrams revealed that the transcriptional response at one dpi was largely distinct, while five- and ten- dpi shared a higher number of DEGs (**Fig. S1A** and **B**), reflecting a transition from initial recognition to more sustained responses. Based on both cytological and transcriptomic evidence, one and five dpi were selected for single-nucleus RNA-seq to capture the early and established stages of host–pathogen interaction.

**Figure 1.**
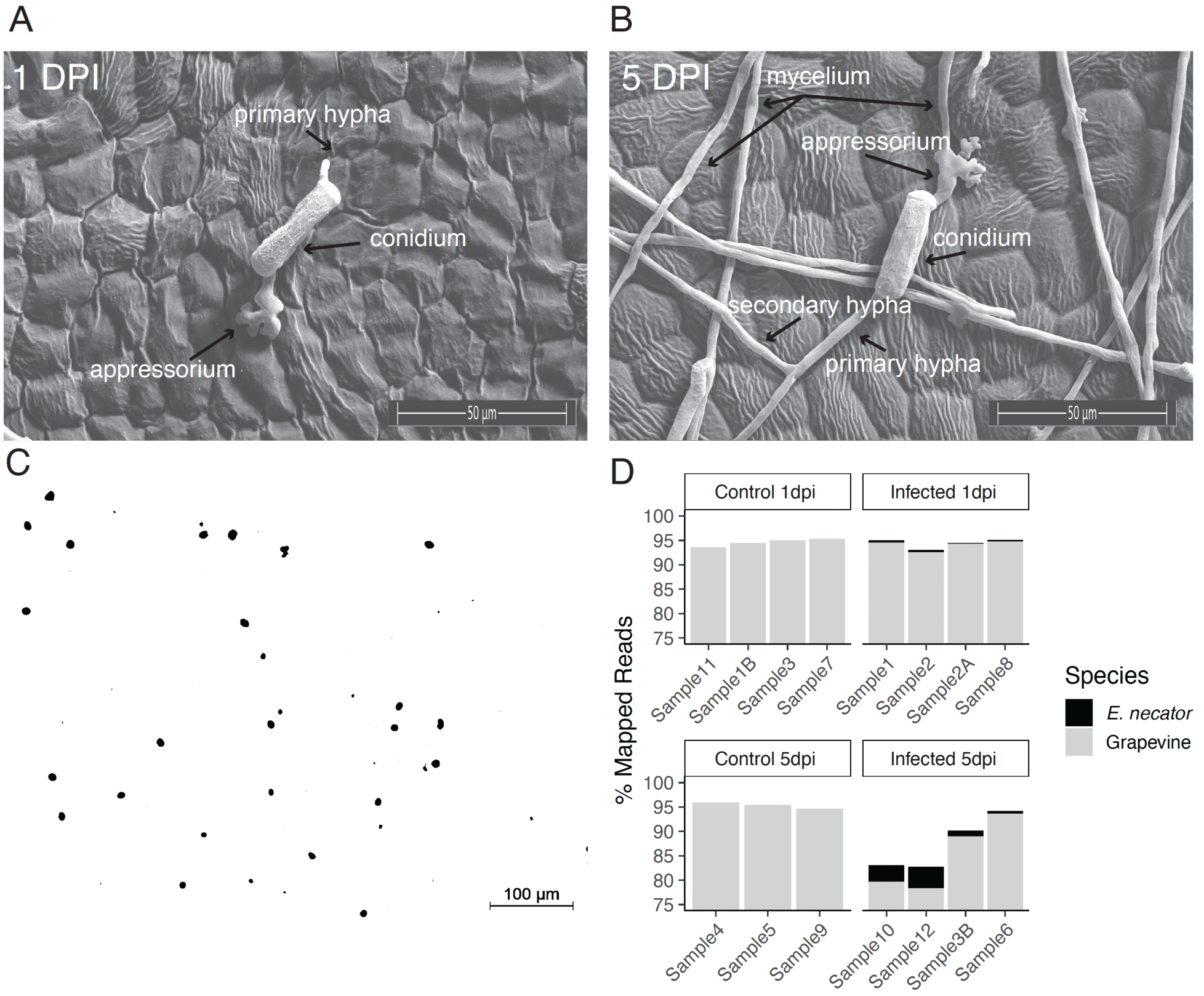
Powdery mildew infection phenology on Cabernet Sauvignon leaves and experimental design. (A) *Erysiphe necator* conidia germinating on Cabernet Sauvignon leaf at one day post infection (dpi). (B) Powdery mildew development on a grapevine leaf at five dpi. (C) Negative field image of DAPI stained nuclei shows intact nuclei membranes and low debris, indicating isolated nuclei passed the quality control for 10x Genomics droplet single-nucleus transcriptomics. (D) Percentage of reads mapping to the Cabernet Sauvignon genome (grey) and the *E. necator* genome (black). Three samples per treatment and dpi were used for grapevine snRNA-seq, and four samples from the infected treatment at respectively one- and five- dpi were used for the *E.* necator snRNA-seq.

High-quality, intact and viable nuclei were isolated from both infected and mock-treated leaves (>1,000 nuclei μl^-1^ in 50 μl; **Fig. 1C**), using two leaves per biological replicate and three biological replicates per treatment and time point. Single-nucleus transcriptomes were generated using the 10x Genomics platform. Following quality control of raw sequencing data, we retained a total of 101,976 nuclei (13,542 ± 4,785 nuclei per sample) including 33,011 and 34,185 nuclei from control and infected samples at one dpi, and 15,850 and 18,930 nuclei at five dpi, respectively (**Table S1**). Samples had on average 606.38 ± 207.28 reads and 423.56 ± 153.23 genes per nucleus (**Table S1**). Control and infected samples had respectively 0 ± 0 % and 0.35 ± 0.15 % at one dpi and 0 ± 0 % and 2.37 ± 1.82 % at five dpi reads mapped to *E. necator* genome (**Fig. 1D**) confirming successful dual-organism profiling.

### Identification of grapevine leaf cell types

After removing pathogen-derived nuclei and data integration, non-linear dimensional reduction using UMAP on the merged samples datasets showed that clustering was primarily driven by cell type identity (**Fig. 2A** and **B**), rather than by individual samples (**Fig. S2A**) or treatment conditions (**Fig. S2B**). Louvain-based clustering identified 21 distinct clusters. To assign cell type identities, we analyzed the expression pattern of a list of previously known leaf cell type markers (**Table S3**; (Baumgart et al., 2025) (**Fig. 2B**). A total of 4,409 cell type markers were found using the FindMarkers function from the Seurat package in R and the results were filtered by adjusted *p* value < 0.01 and log2 fold-change > 1 (**Table S4**). Key marker genes used to define the main cell types are visualized in **Fig. 2B**.

**Figure 2.**
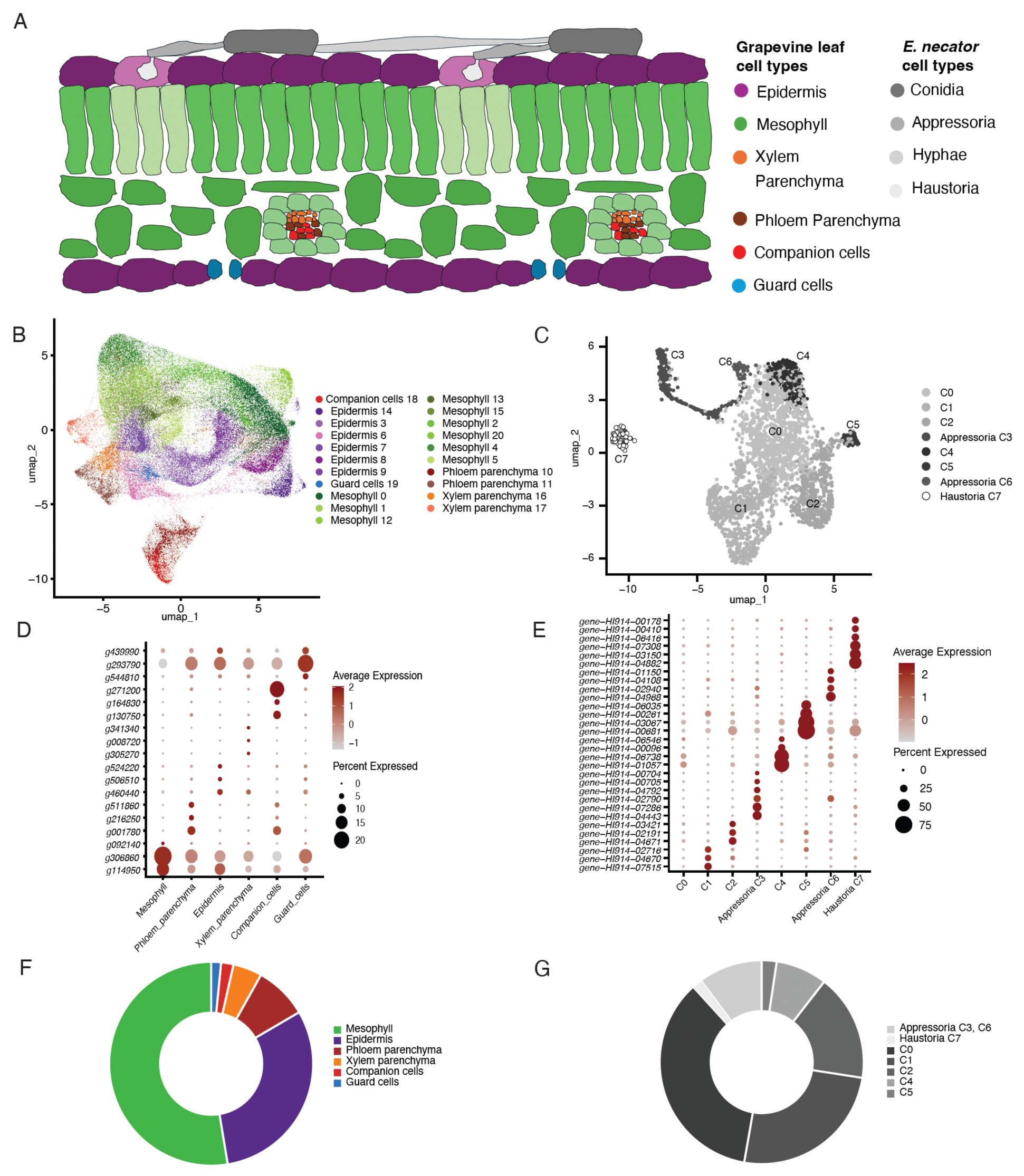
Cluster annotation and cell type marker identification. (A) Schematic representation of the interaction between a grapevine leaf and *Erysiphe necator*. (B) UMAP-based clustering and gene ontology-informed annotation of grapevine leaf cell types, based on single-nucleus RNA-seq from three biological replicates of control and infected leaves at one- and five- dpi. (C) UMAP-based clustering of the *E. necator* nuclei from actively infected grapevine leaves. (D) Percentage of nuclei and scaled average expression levels (mean=0 and variance=1 within each gene using DotPlot function with parameter scale =TRUE) of marker genes for grapevine cell types. (E) Percentage of nuclei per cluster and scaled average expression of marker genes for *E. necator* cell types. (F) Pie chart indicating the relative abundance of each grapevine cell type relative to the total number of nuclei for each grapevine leaf cell type. (G) Pie charts showing the distribution of *E. necator* nuclei across identified fungal cell types and clusters.

The mesophyll was identified in clusters 0, 1, 2, 4, 5, 12, 13, 15, and 20, which showed high expression of photosynthesis-related genes, including transketolase genes (*g533770* and *g238660*), and Calvin cycle-related genes (*g306860;* **Fig 2D**). Epidermal cells were associated with clusters 3, 6, 7, 8, 9, and 14, based on the expression of cuticle biosynthesis genes such as O-acyltransferase WSD1-like (*g524210* and *g524220*) and 3-ketoacyl-CoA synthase 6 (*g506510*) genes (**Fig. 2D**). Phloem parenchyma cells were identified in clusters 10 and 11 by high expression of *PHLOEM PROTEIN 2-LIKE A9-like* genes (*g298430* and *g001780;* **Fig. 2D**), while xylem parenchyma was localized to clusters 16 and 17 based on *WAT1-related protein* genes (*g008720* and *g305270;* **Fig. 2D**). Companion cells were represented in cluster 18, marked by high expression of a sucrose transporter gene (*g271200;* **Fig. 2D**a, and the zinc finger transcription factor *DOF3.1* gene (*g130750;* **Fig. 2D**). Guard cells (cluster 19) were identified based on the guard-cell-specific marker gene *ASPARTIC PROTEASE IN GUARD CELL 1-like* gene *(g439990* and *g544810*), and the 9-cis-epoxycarotenoid dioxygenase *NCED3* (*g293790;* **Fig. 2D**).

Based on this annotation, mesophyll cells accounted for 52.60% of all nuclei and 33,500 expressed genes, followed by epidermis (30.82% nuclei and 33,323 genes), phloem parenchyma (8.44% nuclei and 31,926 genes), xylem parenchyma (4.61% nuclei and 30,127 genes), companion cells (1.96% nuclei and 28,091 genes), and guard cells (1.56% nuclei and 26,091 genes) (**Fig. 2F**). The clear separation of clusters and the assignment of canonical markers to most of them validate the approach and establish a solid foundation for subsequent analyses of tissue-specific gene functions and their dynamics during PM infection.

### Grapevine defense-related genes are expressed in a cell type-specific pattern

After identifying the major cell types in grapevine leaves, we analyzed the cell type-specific expression of gene sets associated with key defense-related pathways. We calculated the number of expressed genes for each set in each nuclei cluster to identify cell type-specificity of defense-related genes. Except for nitric oxide production-related genes, which showed a uniform expression pattern among all cell types (**Table S5**), we found 4,134 defense-related genes expressed in the epidermis, 4,215 in the mesophyll, followed by phloem (3,284) and xylem (2,758) parenchyma, and lastly by companion cells (2,242) and guard cells (1,829; **Table S5**). Genes encoding receptors (NLRs and RLKs), pathogenesis-related genes, and phytoalexin biosynthesis genes were present in almost double percentage in epidermis and mesophyll compared to companion and guard cells, regardless of the PM treatment (**Table S5**). This result is consistent with previous findings in single-cell (Chhillar et al., 2025a) and bulk transcriptomic approaches (Chhillar et al., 2025b).

To further examine these patterns, we visualized cell type-specific expression pattern of defense-related genes using UMAP-based plots. We calculated module scores for each defense-related gene set and the 95% confidence intervals (CI) of the estimated marginal means per each cluster and treatment, enabling us to detect the spatial and cell type-specificity of distinct defense responses. The expression pattern of NLR genes was cluster-specific, although it did not change significantly in response to the PM infection. The NLRs were highly expressed in clusters 8, 9 and 14 of the epidermis, and 0, 20, 8, 12 of the mesophyll (**Fig. 3A and E**), and it was similar to the expression pattern of MAPK-signaling genes (**Fig. 3B and F**). The expression of RLK genes peaked at five dpi in the infected clusters 7 and 9 of the epidermis and 16 of the xylem parenchyma (**Fig. 3C** and **G**). Our data showed that pathogenesis-related genes were more expressed in cluster 7 of the epidermis in response to the PM infection at one and five dpi (**Fig. 3D** and **H**). The cell wall reinforcement-related genes were mostly expressed in vascular tissue regardless of the PM infection (**Fig. S3A**). High average expression of phytoalexin biosynthesis related genes was detected at one dpi in the infected samples, with the highest increase in cluster 7 (Epidermis; **Fig. S3B**). At five dpi, genes related to ROS production were more highly expressed in response to PM in all grapevine leaf tissues, with a significant increase compared to their controls in cluster 7 and 9 of epidermis, 11 of phloem parenchyma, 16 of xylem parenchyma and 13 of mesophyll (**Fig. S3C**). Calcium signaling-related genes were more highly expressed in infected samples at five dpi, especially in cluster 7 of the epidermis and in guard cells (**Fig. S3D**). Infected cluster 7 of the epidermis and guard cells showed an increase in the expression of genes involved in abscisic acid (ABA)-mediated signaling pathway at five dpi (**Fig. S3E**), whereas the expression of genes of the ethylene-mediated signaling pathway decreased in the same clusters (**Fig. S3F**), likely due to antagonism between ABA- and ethylene-mediated signaling pathways. The expression of genes belonging to the jasmonic acid-mediated signaling pathway increased in response to PM at five dpi in cluster 20 of the mesophyll (**Fig. S3G**). Genes involved in the salicylic acid-mediated signaling pathway were highly expressed in the clusters 2, 13, and 15 of the mesophyll and cluster 7 made of the epidermis of the PM-infected samples (**Fig. S3H**).

**Figure 3.**
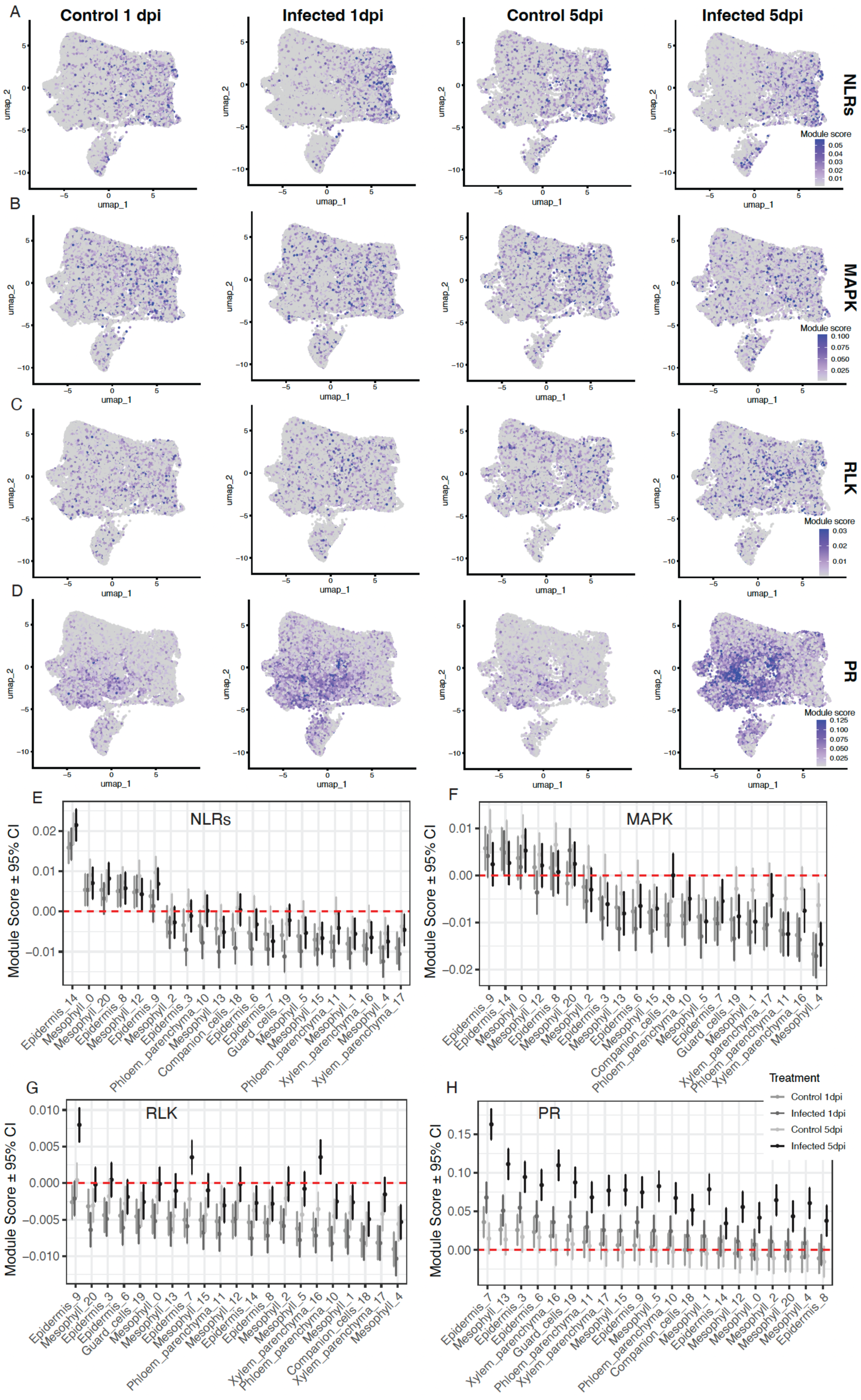
Host defense-related gene expression patterns during powdery mildew infection. (A-D) UMAP-based visualization of average expression per nucleus for defense-related gene sets, calculated using the AddModuleScore in Seurat, across treatment groups (control and infected leaves at one- and five- days post infection, three biological replicates). Panels show the following gene sets: (A) nucleotide-binding leucine-rich repeat proteins (NLRs), (B) mitogen-active protein kinase (MAPK), (C) receptor-like kinase (RLK), (D) pathogenesis-related (PR). (E-H) Module score estimated marginal means 95% confidence intervals (CI) for the same gene sets: (E) PR genes, (F) cell wall reinforcement, (G) phytoalexin biosynthesis, and (H) reactive oxygen species production. A positive module score indicates a higher average expression than expected whereas a negative module score indicates a lower average expression than expected, given the average expression of this module across the whole dataset.

### Pseudobulk differential expression analysis reveals grapevine leaf cell type-specific responses to powdery mildew

With cell types and defense-related gene expression patterns defined, we next examined how each cluster responded transcriptionally to *E. necator* infection. To identify DEGs across clusters and time points, we performed differential expression analysis using a pseudobulk approach (Tenorio Berrío et al., 2025), followed by gene ontology enrichment analysis. Because snRNA-seq generates data from thousands of nuclei per sample, treating individual nuclei as replicates would underestimate biological variance and inflate statistical significance. The pseudobulk method overcomes this by aggregating counts per cluster within each biological replicate, allowing for variance estimation across true replicates and improving statistical power (Junttila et al., 2022; Squair et al., 2021). Of the DEGs, 183 and 754 were in common with the bulk-tissue RNA-seq at respectively one and five dpi (**Fig. S1C**).

At one dpi, the most transcriptionally affected clusters were 3 and 9 of the epidermis, with respectively 65 and 28 downregulated genes and 104 and 133 upregulated genes, respectively, followed by cluster 2 of the mesophyll, with 15 downregulated and 81 upregulated genes (**Fig. 4A** and **B**). From a transcriptional point of view, other clusters were only marginally affected by the PM infection (**Fig. 4A** and **B**). By five dpi, transcriptional reprogramming expanded across tissues. The cluster 0 of the mesophyll showed the highest number of both down- (266) and up-regulated genes (339), indicating an active role of the mesophyll in PM response, at least in susceptibility-type response (**Fig. 4C** and **D**). Clusters 7 and 9 were the most affected clusters of the epidermis with respectively 202 and 243 downregulated genes and 239 and 337 upregulated genes, respectively (**Fig. 4C** and **D**).

**Figure 4.**
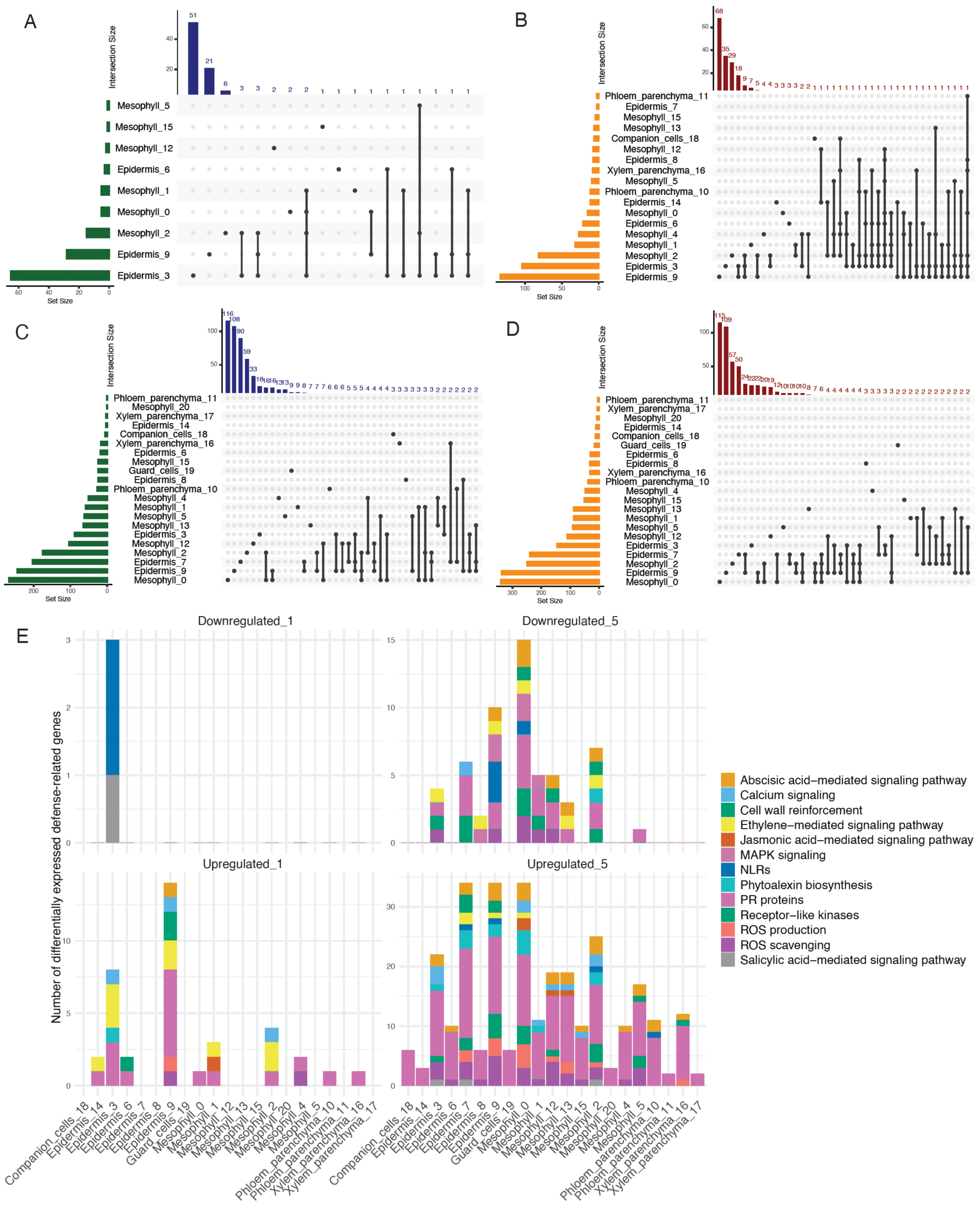
Grapevine pseudobulk snRNA-seq analysis identified both shared and cell-type-specific differentially expressed genes. (A-D) Comparison of downregulated (A, C) and upregulated (B, D) genes in response to PM between clusters at one dpi (A, B) and five dpi (C, D). (E, F) Gene ontology of down- (E) and upregulated genes (F) at one and five dpi in infected leaves compared to matched controls.

These transcriptional patterns were further supported by functional annotation of DEGs by cell type and time point (**Table S6**). At one dpi, the most responsive cluster was cluster 9 of the epidermis, followed by cluster 3 (**Fig. 4E**). The cluster 3 of the epidermis showed a downregulation of two NLR genes (*g475560* and *g526350*), and one salicylic-acid related gene (*g196080*). At five dpi, cluster 7 and 9 of the epidermis and cluster 0 of the mesophyll were the most responsive to the PM infection (**Fig. 4E**).

We noticed the expression suppression of genes associated with thiamine metabolic process in cluster 9 of the epidermis, which can indicate the pathogen capacity to reduce the expression of pathogenesis-related (PR) genes, enhance callose deposition at infected sites, and lead the accumulation of salicylic acid, a key hormone in systemic acquired resistance (**Fig. S4**). The functional annotation of the DEGs in cluster 2 of the mesophyll indicated an expression decrease of genes involved in catabolic processes, including defense catabolic enzymes (endochitinase, *g369980*; glucan endo1,3-beta-glucosidase, *g088060*), likely induced by pathogen effectors. The cluster 14 of the epidermis, rich in NLR genes (**Fig. 3A**) showed enriched nucleotide-related metabolic processes among its DEGs in response to PM (**Fig. S4**; **Table S6**), which indicate an activation of NLRs (e.g., ATB-bound NLRs are active, whereas ADP-bound NLRs are inactive; (Wang et al., 2019a). At one dpi, the increased expression of genes belonging to the terpene, isoprenoid and hydrocarbon metabolic processes in companion cells (**Fig. S4; Table S6**) suggests an active participation of this cell type at early-stage infection response to *E. necator*.

At five dpi, the biological process of defense response to fungus was enriched among the DEGs of most clusters of the epidermis, mesophyll, phloem and xylem parenchyma, guard cells and companion cells (**Fig. S4; Table S6**). We noticed an expression suppression of genes involved in actin filament organization in the highly responsive cluster 7 (**Fig. 3D)** of the PM-infected epidermis, which could indicate direct cellular defense or pathogen effectors disrupting actin. Genes associated with lipid transport was also strongly downregulated in cluster 9 of the epidermis in response to PM (**Fig. S4; Table S6**). Genes from the phospholipid and glycerophospholipid metabolic process were more lowly expressed in response to PM in cluster 2 of the mesophyll. The most enriched biological processes among the DEGs at five dpi were in cluster 9 of the epidermis and included receptor-like genes (*g175930* and *g162640*), PR genes (*g206640* and *g502040*), and MLO-like genes (*g189080* and *g484430*) (**Table S6**).

### Gene co-expression analysis reveals cell type-specific hubs and modules involved in powdery mildew response

To further investigate the regulatory architecture underlying cell type-specific responses to PM, we performed hdWGCNA. We constructed the gene network based on epidermal cells and calculated the module eigegenes (ME) for each module using the whole dataset to interrogate the cell-type specificity of these modules’ expression programs across all cell types (Morabito et al., 2023). During network construction, a soft threshold power (β) of 8 achieved a scale-free topology fit index of 0.90 (**Fig. S5A**). This analysis identified seven gene modules related to the epidermis (**Fig. 5**), with unassigned genes grouped in the grey module. Hierarchical clustering based on expression patterns organized the modules into three groups: M1 and M7; M4, M5, M2, and M6; and M3 alone (**Fig. 5A**). Relationships among modules were further supported by gene network topology (**Fig. 5B**). Genes with the highest module membership scores, referred to as hub genes, are shown in **Fig. S5B**. Module-specific gene expression and the spatial distribution of hub genes are visualized in UMAP plots (**Fig. 5C** and **D**).

**Figure 5.**
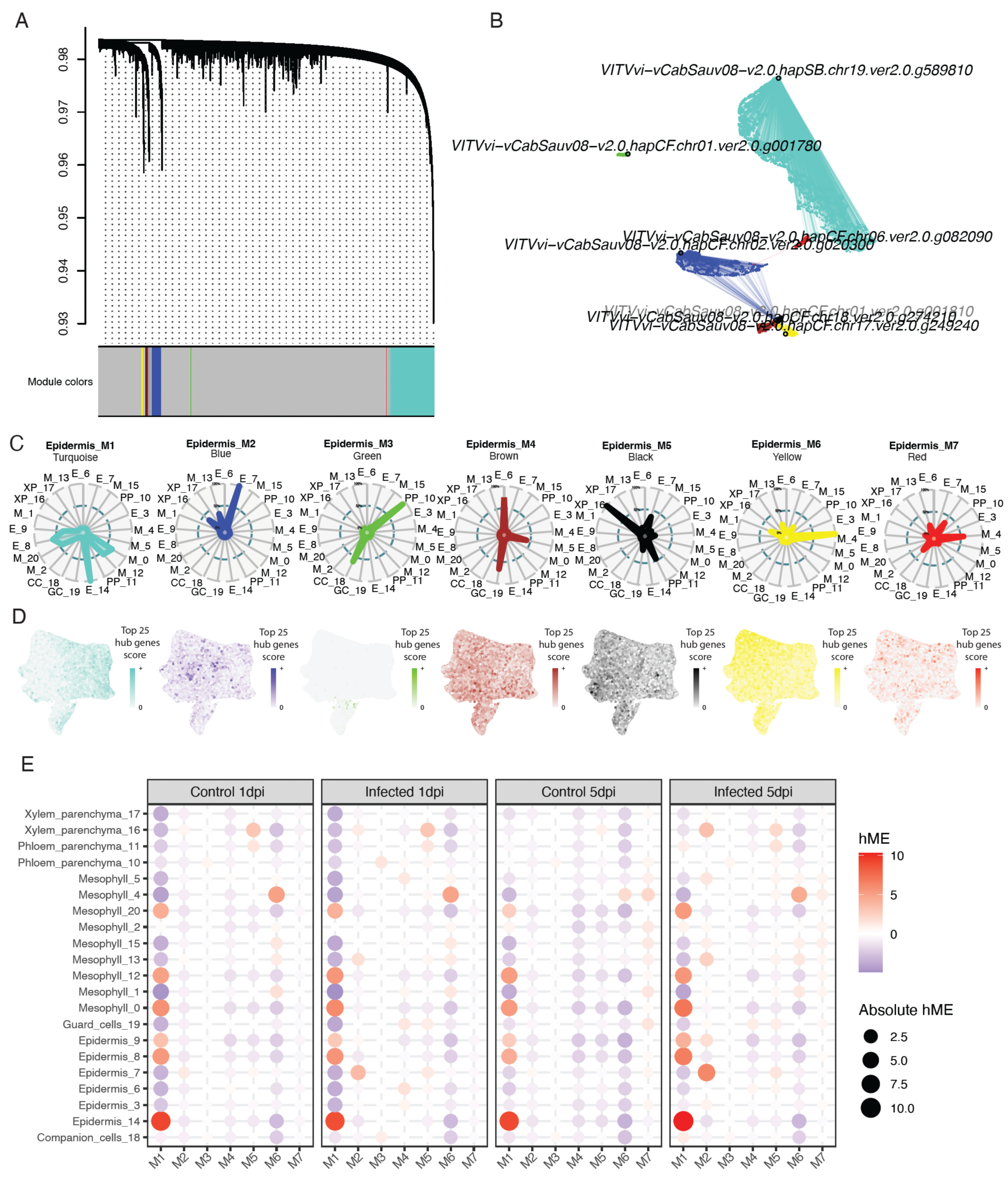
Epidermis-related module detection and eigengene expression patterns from hdWGCNA analysis of grapevine single-nucleus RNAseq data. (A) Hierarchical clustering dendrogram of grapevine genes based on topological overlap, with epidermis-related modules identified by dynamic tree cutting and assigned distinct colors. (B) Network visualization of representative hub genes and their interactions with selected epidermis-related modules. (C) Radar plots showing the relative expression level of epidermis-related co-expression modules in each cluster (E=”Epidermis”, M=”Mesophyll”, PC=”Phloem parenchyma”, XP=”Xylem parenchyma”, CC=”Companion cells”, GC=”Guard cells”). Concentric circles represent 100% (most external circle), 50% (blue circle) and 0% (center of the radar plot). (D) UMAP plots displaying top 25 hub genes UCell score for each module. (E) Average hME scores across cell type clusters and treatments. Facets correspond to treatment groups (control and infected at one and five dpi), allowing comparison of module activity over the course of infection.

The module M1 had high hME levels in multiple clusters, including the epidermis clusters 8, 9, and 14, and the mesophyll clusters 0, 13, and 20, regardless of the PM treatment (**Fig. 5E**). Its hub genes were stress related genes (**Table S7; Fig S5B**), such as DUF594 family protein-coding gene (*g589810*), disease resistance genes (*g573170*, *g163360, g163000, g589850, g458780*), vacuolar protein-sorting proteins (*g023470, g320020, g060980*), suggesting a general stress response independent of PM infection. The high percentage of NLR and MAPK genes in this module’s hubs suggests the involvement of this module in ETI response (**Table S8**).

The module M2 was associated with PM induced responses. It showed high hME levels in infected samples at one dpi in the clusters 7 (Epidermis), 13 (Mesophyll) and 16 (Xylem Parenchyma), and at five dpi in the epidermis clusters 7 and 9, the mesophyll clusters 13, 5, and 15, and the cluster 16 made of xylem parenchyma (**Fig. 5E**). This module’s hub genes were composed of PR genes (**Table S7; Fig S5B**), including genes encoding thaumatin-like proteins (*g020300, g020290*), major allergen Pru ar 1-like (*g059230, g059220, g355410*), endochitinase EP3-like (*g369980, g074560, g074570*), dirigent proteins (*g372250, g250530, g352800*). The high percentage of RLK and PR genes in this module’s hubs suggests the involvement of this module in PTI response (**Table S8**).

The module M3 was strongly expressed in the cluster 10 made of phloem cells and cluster 18 of companion cells (**Fig. 5E)** and showed hub genes related to phloem and sieve (**Table S7; Fig S5B**; e.g., *g001780, g298430, g006660*), supporting the accuracy of the cluster annotation.

The module M4 showed high hME average in the cluster 19 made of PM-infected guard cells (**Fig. 5E)** and had hub genes related to ABA-mediated signaling pathway (**Table S7; Fig S5B**), including abscisic stress-ripening protein 2-like genes (*g274210, g569270*) and 9-cis-epoxycarotenoid dioxygenase *NCED3* (*g293790, g588250*). This module had higher hME average in PM-infected samples in the cluster 19 (GuardCells) at both one- and five- dpi, and in the clusters 6 (Epidermis) and 5 (Mesophyll) at one dpi, suggesting a potential role of the ABA-mediated signaling pathway in PM response. This is further supported by the high module membership (kME; i.e., Pearson correlation between a gene’s expression profile and the eigengene of its module) of major latex protein (MLP)-like proteins within this module (e.g., *g005040, g301630;* **Table S7**).

The module M5 showed elevated hME in the clusters 16 (Xylem Parenchyma) and 11 (Phloem Parenchyma) under all conditions and in the clusters 7 (Epidermis) and 19 (Guard Cells) in PM-infected samples at one and five dpi (**Fig. 5C** and **E**). This module’s hub genes contained ribosomal related protein-encoding genes (**Table S7; Fig S5B**), suggesting an increased protein synthesis, possibly reflecting tissue recovery or active growth (e.g., in protoxylem and protophloem).

The module M6 was the most strongly expressed in mesophyll clusters and showed a high number of photosynthesis-related hub genes (**Table S7; Fig S5B**), confirming cluster annotation (**Fig. 5C, D** and **E**).

The module M7 showed elevated hME in mesophyll clusters at five dpi, independently of the treatment (**Fig. 5E**) and high number of hub genes involved in cell wall reinforcement (e.g., S-adenosylmethionine decarboxylase, exordium-like proteins), photosynthesis (e.g., Thylakoid Soluble Phosphoprotein TSP9, Rubisco activase), and redox regulation (e.g., ferredoxin, glutaredoxin domain-containing cysteine-rich protein 1, copper transporter 6-like and catalase isozyme 1) (**Table S7; Fig S5B**). Interestingly, module M7 connected module M1, associated to high NLR gene expression and ETI, with module M2, associated to high RLK gene expression RLK and PTI (**Fig. 5B**). Hub genes in M7 included, *MAPK15* (signal transducer), *wall-associated receptor kinase-like 16*, *NDR1-like* (plasma membrane-linked signal regulator produced downstream of RLK signals and critical for CNL-type NLRs; (Knepper et al., 2011; Wang et al., 2023), transcription factors *MKY1R1* and *TCP7* (**Table S7**), which likely regulate and bridge PTI and ETI response.

### Single-nucleus expression atlas of *Erysiphe necator* infection structures during powdery mildew development

Leveraging the dual snRNA-seq approach, we extracted the sequencing data corresponding to *E. necator* from the PM-infected samples and performed unsupervised clustering to define transcriptionally distinct fungal cell types during grapevine infection. This analysis enabled us to map gene expression associated with key infection structures, including appressoria, haustoria, and hyphae at various developmental stages, providing a framework to link fungal morphology with function during pathogenesis.

After selecting fungal reads and applying log-normalization and SCTransform, Louvain-based clustering identified eight distinct fungal clusters (**Fig. 2C**). Clusters C3 and C7 showed clear separation from the others, and only half of the clusters (C0, C1, C3, and C7) included nuclei from one dpi, suggesting that these clusters represent early stages of infection. Clusters were identified based on genes expressed in at least 10% of nuclei within a given cluster. Cluster C7 was identified as haustoria based on the expression of two hallmark gene families previously described in PM pathogens: Candidate Secreted Effector Proteins (CSEPs; Spanu et al., 2010) and RNase-Like Proteins Associated with Haustoria (RALPHs; Pedersen et al., 2012; Pennington et al., 2019; Spanu, 2017). A total of 100 CSEPs were consistently expressed, of which 57 were specific to cluster C7. Similarly, 71 putative RALPHs encoded by the *E. necator* genome, 43 were consistently detected, with 40 exclusively expressed in cluster C7 (**Table S9** and **S10**).

Clusters C3 and C6 were identified as appressoria. These clusters showed strong expression of genes coding for *Egh16*-like virulence factor domains, with homologs in many pathogenic fungi, including *Magnaporthe grisea* GAS2/MAS1 proteins, which are specific to the appressoria (Grell et al., 2003; Xue et al., 2002). Of the 11 *Egh16*-like genes in the genome, six were consistently detected, four of which were exclusively and highly expressed in clusters C3 or C6 (**Table S9** and **S10**). Clusters C0, C1, C2, C4, and C5 showed more heterogeneous expression profiles, likely representing different hyphal developmental stages. Some of these likely represent hyphal stages, including actively growing hyphae, mature hyphae, or hyphae preparing for conidiophore formation. However, due to the lack of definitive hypha-specific markers, these clusters were annotated numerically and are referred to as hyphal clusters. A summary representation of the total nuclei per cluster or cell type is presented in **Fig. 2G**.

In total, we detected the expression of 6,544 of the 7,146 predicted *E. necator* genes. For downstream analysis, we focused on 1,760 genes (24.6%) that were expressed in at least 10% of nuclei within a given cluster (**Table S9** and **S10**). On average, each cluster consistently expressed 676 ± 36 genes, with the haustoria cluster C7 among the top three in total expressed genes (**Table S11A**). To further explore the transcriptional profile of the identified cellular states, we examined cluster-specific gene expression profiles. A total of 738 genes were uniquely expressed in a single cluster, reflecting distinct molecular programs across fungal structures (**Table S11B**). Of these 738 genes, 248 were expressed exclusively in the haustoria cluster C7, 119 in the appressorial cluster C3, and 102 in the appressorial cluster C6 (**Table S11C**). It is notable that at one dpi, we detected expression of 73%, 94%, and 0% of the uniquely expressed genes in the haustoria cluster C7, appressorial cluster C3, and appressorial cluster C6, respectively. In contrast, hyphal clusters (e.g., clusters C0, C2, and C4) contained the fewest uniquely expressed genes, suggesting more transcriptionally conserved or general roles.

### Gene co-expression analysis reveals cell-specific hubs and modules associated with appressoria and haustoria clusters

To investigate the transcriptional programs associated with the development of haustoria and appressoria, we applied hdWGCNA to identify co-expression modules in fungal single-nucleus data. This analysis identified 26 co-expressed modules (**Table S12; Fig. S6A**), 10 of which showed high levels of expression in haustoria or appressoria cluster (**Fig. S6B**). Module-specific gene expression patterns are visualized in the UMAP and dot plots of **Fig. S6C** and **D**. To keep consistency in the analyses, the next paragraphs are focused on the 1,760 genes expressed in at least 10% of nuclei within a given cluster (**Table S10**).

Modules fM2 and fM7 were high hME in the haustorial cluster C7 (**Fig. S6 B** to **D**), with fM2 representing the largest module. As expected, fM2 included 59 CSEPs and 39 RALPHs (**Table S10**). Additional genes in this module included: four chitin synthases, associated with fungal cell wall production or modification; seven histone-related proteins, associated with modulation of gene expression; and various cell cycle-related proteins, likely associated with gene expression regulation, membrane and cell wall production.

Appressorial cluster C3 was characterized by an elevated hME of modules fM3, fM20, fM23, and fM24 (**Fig. S6B** to **D**). For instance, the module fM3 contained genes encoding two cutinases, two MARVEL domain proteins associated with membrane interaction (Douglas et al., 2013; Peng et al., 2024), two Egh16-like virulence factors, and two vacuolar transporters. Module fM23 included a hydroperoxidase I/catalase and a p450 (*CYP52*) genes, while module fM24 grouped genes coding for a NADPH oxidase (NoX), two peptidases, two additional Egh16-like virulence factors, and two vacuolar transporters (**Table S10**).

Appressorial cluster C6 presented high hME of modules fM4 and fM5 (**Fig. S6B** to **D**). These modules included additional Egh16-like virulence factor genes, as well as genes encoding glucanosyltransferases involved in cell wall remodeling and integrity, plasma membrane fusion protein *PRM1* associated with cell fusion (Aguilar et al., 2007; Heiman and Walter, 2000) and five ubiquitin-related proteins (**Table S10**). Finally, modules fM8 and fM11 presented an elevated hME in both appressorial clusters (C3 and C6) and included genes coding for a serine/threonine-protein kinase related to cell wall degradation, appressorium formation, a Myb-like DNA-binding domain protein associated with gene regulation, and a protein *PAL1* involved in cell morphology (Chen et al., 2022)

### Potential new markers of *Erysiphe necator* infection structures during powdery mildew development

The distinct transcriptional profiles of each *E. necator* clusters enabled the identification of novel candidate markers for specific infection structures (**Fig. 2E**; **Fig. S7; Table S13**). Among these, members of the CE5 CAZyme family, which encode cutinases, emerged as promising markers for appressoria identity (*gene-HI914-00705*, *gene-HI914-00704*, *gene-HI914-04108*). Of the five predicted CE5 in the *E. necator* genome, three were expressed, all found in appressorial clusters (two in C3 and one in C6), consistent with their expected role in cuticle degradation during host surface penetration (Arya and Cohen, 2022; Novy et al., 2021).

Additional genes uniquely and highly expressed in appressorial cluster C3 included those encoding proteins with appressorial penetration-associated domains (*gene-HI914-04792*; implicated in host invasion), ferritin-like domains (*gene-HI914-04443*; potentially involved in iron storage and oxidative stress regulation), and MARVEL domain-containing proteins (*gene-HI914-02400, gene-HI914-07576*; associated with membrane trafficking and polarity during appressorium development; **Fig. S7; Table S13**). Cluster C6 showed specific expression of genes encoding a plasma membrane fusion protein (PRM1; *gene-HI914-01150*), known to mediate membrane merging during cell-cell or cell-host interactions, AB hydrolase domain-containing proteins (*gene-HI914-02940)* usually associated with plant cell wall component degradation and proteins with double-stranded RNA-binding motifs, which may participate in post-transcriptional regulation or host RNA silencing suppression.

The haustorial cluster (cluster C7) expressed all three NAD nucleotidase genes encoded in the *E. necator* genome, with two (*gene-HI914-06416*, *gene-HI914-00410*) showing high expression levels, potentially enabling NAD turnover at the host–pathogen interface. Other genes expressed in this cluster included those with putative roles in nutrient uptake and metabolic coordination, such as sugar porter family proteins (*gene-HI914-03150*; carbohydrate import), cytochrome P450s (*gene-HI914-00178*; CYP53; involved in wax degradation and secondary metabolism), aminotransferases (*gene-HI914-07308*, *gene-HI914-03062*; nitrogen assimilation), dynamin-GTPases (*gene-HI914-02306*; membrane remodeling), and uridine kinases (*gene-HI914-04033*; nucleotide salvage pathways), among others (**Fig. 2E**; **Fig. S7; Table S13)**.

Other clusters also showed distinct marker gene expression. For example, in cluster C1, GH132 (*gene-HI914-07515*) with glucan glucosidase functions (fungal cell wall remodeling) and histone H4 genes (*gene-HI914-04670*; associated with DNA replication and cell cycle) were highly expressed (**Fig. 2E**; **Fig. S7; Table S13)**. Cluster C2 was characterized by expression of GH13+CBM48 (gene-HI914-02191), a glycoside hydrolase with glycogen-binding function, and a thioesterase-like superfamily member (*gene-HI914-03421*). In cluster C4, candidate markers included a type 1 polyketide synthase (*gene-HI914-06546*), 3−dehydroquinate synthase (*gene-HI914-01057*; involved in amino acid biosynthesis), and glutaredoxin (*gene-HI914-00096*; associated with iron homeostasis). Finally, cluster C5 showed specific expression of genes encoding a cell wall mannoprotein (*gene-HI914-00681*; cell wall structure), perilipin protein (*gene-HI914-03067*; lipid storage), and ABC-2 type transporters (*gene-HI914-06035*; stress tolerance). Additional experiments will be necessary to definitively associate these transcriptomic profiles with specific fungal hyphal stages and confirm their utility as structural markers.

## Discussion

This study presents the first application of single-nucleus transcriptomics to the study of the infection between grapevine and *E. necator* during the development of powdery mildew. Beyond demonstrating the power of this approach to resolve cell type-specific transcriptional dynamics, we provide optimized protocols for nuclei isolation, a reproducible analysis pipeline, and a list of over 4,000 novel cell-type gene markers (**Table S3**) that will facilitate future single-cell studies in grapes.

Our single-nucleus gene expression atlas, generated from over 100,000 nuclei across 12 independent samples, captured all major cell types in grapevine leaves. We identified epidermal cells by the expression of cuticle biosynthetic genes, mesophyll by photosynthesis-associated genes, phloem, xylem parenchyma, and companion cells using previously reported markers (Baumgart et al., 2025), and guard cells based on genes specifically expressed in this cell type (e.g., *ASPARTIC PROTEASE IN GUARD CELL 1-like*; Yao et al., 2012). Accurate cell type annotation was essential for characterizing PM responses across cell types and for detecting cellular response heterogeneity. The mining for robust cell type markers is a critical step for implementing single-cell transcriptomics in non-model plant systems.

Although much is known about plant responses to PMs (Qiu et al., 2015), this study provides seminal evidence of cell type-specific responses to PM and reveals a complex network of signaling and defense response across distinct cell types. For example, we detected a higher proportion of defense related genes expression in the epidermis and mesophyll, particularly for RLK, NLRs, and PR related genes. A similar pattern was also observed in *Arabidopsis* in response to *Pseudomonas syringae* DC3000 (Delannoy et al., 2023), suggesting that vascular and guard cells may rely on distinct immune response systems. However, other studies have shown higher NLR expression in vascular cells (Tang et al., 2023). The detection of infected vascular cells remains challenging, due to the lower abundance in leaf tissue (Delannoy et al., 2023), which may explain inconsistencies across studies. Further research is needed to clarify the differences in innate immune capacity among epidermal, mesophyll, and vascular cells, ideally also by examining responses to vascular-colonizing pathogens, such as *Xylella fastidiosa* (Rapicavoli et al., 2018).

At the early stage of the PM infection, we observed epidermis-specific overexpression of defense related genes (**Fig. 3C** and **D**), identifying nuclei in close proximity to fungal structures and confirming a strong correlation between cellular response and adjacency to the invading pathogen (Tang et al., 2023; Zhu et al., 2023). In parallel, also at one dpi, a small subset of mesophyll cells also exhibited immune response activation and signaling activity (**Fig. 4A** and **B**), suggesting an early contribution of mesophyll cells to the defense response, consistent with findings from other plant-pathogen single-cell studies (Nobori et al., 2025). As PM progressed, the transcriptional response broadened to include both the epidermis and the mesophyll, while vascular tissue and guard cells still showed limited activation (**Fig. 4C** and **D**). The temporal shift in cluster responses reflects the progression of infection: early defense is concentrated in the epidermis, the site of fungal entry and cellular invasion, followed by a secondary response in the mesophyll, indicating the spread of the pathogen’s impact beyond the site of initial colonization. This progression likely reflects coordinated, tissue-specific defense response signaling and metabolic reprogramming during disease development.

The snRNA-seq approach also enabled the detection of spatial heterogeneity within cell-types. Interestingly, among the epidermal clusters, we observed localized expression of RLKs associated with cell surface signaling (**Fig. 3C**), suggesting the activation of PTI, a signal that would have been masked in bulk RNA-seq data. This was further supported by the overexpression of PR genes in the same clusters (**Fig. 3D**). Consistent with this finding, a rare PRIMER cell state was recently described in *Arabidopsis*, located at the nexus of immune-active hotspots (Nobori et al., 2025). These cells act as local immune hubs that activate defense responses and relay signal responses to the surrounding cells during infection.

The NLR genes were expressed in specific clusters of the epidermis and mesophyll (**Fig. 3A**) and were predominantly constitutively expressed (**Fig. 3E**), suggesting that their activation may occur by oligomerization into resistosomes (Hu and Chai, 2023; Huang et al., 2023; Wang et al., 2019b). The spatial and cellular heterogeneity of NLR supports a model of localized activation of ETI, distinct from the spatial pattern of PTI. Recent studies suggest that ETI can be suppressed by the pathogen effectors, but restored in surrounding cells, leading to local acquired resistance (Jacob et al., 2023; Nobori et al., 2025). Our results align with this model, with epidermal nuclei clusters characterized by strong activation of RLK-mediated PTI (**Fig. 3C**), including overexpression of pathogen-related proteins and oxidative stress, but low NLR expression, even slightly below the matched control (**Fig. 3E**). These epidermal clusters likely represent cells in direct contact with the pathogen. In contrast, neighboring epidermal and mesophyll clusters showed ETI signatures mediated by NLRs (**Fig. 3A**), and may represent regions of peripheral immune activation, consistent with localized acquired resistance (Jacob et al., 2023). Gene co-expression network further supports the dual-layered architecture of the plant immune system, as the two major modules (M1 and M2) were associated with PTI and ETI (**Fig. 5B** and **D**). The two module networks were connected by a third module (M7) which allowed to detect candidate genes (**Table S6**) responsible of the crosstalk between the ETI and PTI. Notably, M7 showed high expression in some mesophyll clusters (**Fig. 5C**), indicating an active role for the mesophyll in immune signaling coordination during PM.

This study also provides the first detailed transcriptional atlas of *E. necator* infection structures, revealing structure-specific gene expression programs associated with surface attachment, penetration, nutrient uptake, and immune suppression. Studying the transcriptomes of *E. necator* has been challenging due to the difficulty of isolating sufficient pathogen material from actively infected tissue and the absence of established methods for isolating haustoria of powdery mildew fungi, unlike for rust pathogens (Cantu et al., 2013). As a result, transcriptomic studies of powdery mildew infections have relied on bulk samples limiting the ability to resolve specific fungal structures (Hu and Chai, 2023; Wang et al., 2019b). Here, we demonstrate that single-nucleus transcriptomics can effectively detect virulence factors expressed in key infection structures. Known markers were highly and consistently expressed in three of the eight nuclei clusters, enabling the confident identification of haustoria and appressoria, and validating of the approach. More than 240 genes were uniquely expressed in haustorial nuclei and over 200 in appressorial nuclei, indicating that the mechanisms of surface attachment and penetration (appressoria) and nutrient acquisition and immune suppression (haustoria) involve substantial transcriptional reprogramming.

The CSEPs and RALPHs gene families have been consistently associated with haustorial function, and their strong, specific expression in cluster C7 (**Table S10**) supports its annotation as haustorial nuclei. These families have been extensively studied in other biotrophic pathogens through various approaches, including bulk RNAseq at early infection stages and haustoria-enriched samples (Cantu et al., 2013; Godfrey et al., 2010; Pedersen et al., 2012; Sharma et al., 2019; Spanu, 2017). While members of these families had been previously identified in *E. necator* (Jones et al., 2014; Zaccaron et al., 2023), this is the first confirmation of their haustorial expression at single cell resolution. Moreover, our data revealed that subsets of these gene families are exclusively or predominantly expressed in haustorial clusters, while others are expressed in non-haustorial clusters. These findings suggest that expression of CSEPs is not restricted to haustoria, as previously reported in other powdery mildew species, where haustoria-containing plant epidermis and epiphytic structures were analyzed, and while the majority of CSEPs families were predominantly expressed in the haustoria samples, some families were expressed at similar levels in both kinds of tissues. (Aguilar et al., 2016; Pedersen et al., 2012). This finding provides a foundation for refining and functionally characterizing specific subsets of haustorium markers. In contrast, appressorium-specific markers in powdery mildew fungi have been more difficult to identify, due to the lack of enrichment protocols that isolate appressoria from other epiphytic structures. Consequently, previous studies relied on infection timing or differential expression to infer candidate markers. While genes such as *MAPK*, *clap1*, *cPKA*, *Egh16*-like proteins, and others (Both et al., 2005; Grell et al., 2003) have been proposed as markers of appressorial development, only *Egh16*-like domain genes showed a strong and exclusive association with two clusters in our dataset (**Table S10**). The expression of homologs of other previously suggested appressorial markers, like *MAPK* was detected but lacked specificity or consistent expression in the cluster.

Co-expression modules associated with appressoria further supported the identification of this infection structure (**Fig. S6B** to **D; Table S10)**. Module fM23 included genes such as catalase-peroxidase and *CYP52* (fM23), which are involved in the degradation of plant-derived defense compounds and cuticle waxes (Guo et al., 2019; Liu et al., 2007; Ortiz-Álvarez et al., 2020). Additionally, genes such as a serine/threonine protein kinase and PAL1, also present modules in fM8 and fM11, have been shown to be essential for turgor pressure buildup and penetration in *M. grisea* and *M. oryzae*, respectively (Chen et al., 2022). The new candidate markers proposed for identifying *E. necator* infection structures should enhance the accuracy of cluster annotation in single-cell studies. However, additional experiments are needed to link the specific physiology and development of hyphal cells with the proposed markers.

Together, these results underscore the value of single-nucleus transcriptomics for resolving spatial immune dynamics in a susceptible host that ultimately permits *E. necator* colonization and sporulation. Future studies applying this approach to resistant grapevine genotypes will be critical for dissecting the host and pathogen transcriptional dynamics during effective immune responses. These analyses will help distinguish basal defense mechanisms from those that successfully block infection and, using virulent *E. necator* strains, reveal how the pathogen overcomes resistance. Ultimately, this work will support the strategic deployment of resistance loci, guide breeding efforts toward durable resistance, and provide insight into pathogen strategies to evade host immunity (Paineau et al., 2024).

## Materials and methods

### Biological material and growth conditions

Detached grapevine leaves from Cabernet Sauvignon clone FPS08 were used for this study. The infected leaves were inoculated with the PM *E. necator* strain UCD-TS1, using a modified protocol from Reifschneider and Boiteux, 1988. A 15-day PM-infected *in vitro* leaf was used as inoculum in a closed 30-cm cake box with five sterilized *in vitro* leaves (open Petri dishes). Constant air flow was provided for 1.5 min from the top of the *inoculum* using a wide-end pipette tip attached to a rubber hose attached to an air nozzle outlet. This protocol was optimized to ensure a uniform distribution of conidia density (∼20 conidia per mm^2^) on the leaf surface. Mock leaves were treated in the same way, but no inoculum was present in the closed box. Leaves were placed in a growth chamber, with 16h photoperiod, T ranging from 16C (night) to 23C (day), and humidity at ∼60%. After 24h, the Petri dishes were sealed with parafilm. For scRNA-seq, a total of 15 samples (four for control and PM-infected at one day post infection (dpi); three for control and four for PM-infected at five dpi) were harvested at one and five dpi. For RNA sequencing (RNA-seq), three mock- and PM-inoculated leaves were harvested at one, five and ten dpi at noon. Inoculum and harvesting were done at noon for every experiment. Three biological replicates were included for each assay and each treatment, and time point.

### SEM imaging

For scanning electron microscopy (SEM), mock and PM-infected 5x5-mm leaf sections were immersed in 2.5% glutaraldehyde 2% paraformaldehyde in 0.1M phosphate buffer and kept at 4°C overnight. The leaf sections were dehydrated through ethanol, critical point dried, gold sputter coated, and observed at the SEM.

### Single-nuclei isolation

Single-nuclei isolation from grapevine leaves was optimized using a Plant Nuclei Isolation/Extraction Kit (CelLytic™ PN, Sigma). Two fresh frozen intact leaves (∼1.5 g) were finely chopped in 1 mL Nuclei Isolation Buffer (NIB) for 1 min using a razor blade on a Petri dish. The sample was transferred in a 50-mL Falcon tube, NIB buffer was added to reach 25 mL volume, slowly inverted two times and incubated on ice for 15 min. The sample was filtered using a 20 μm steriflip vacuum filter and then centrifuged at 5000 rpm for 10 min. Nuclei Isolation Buffer-A was added (5 mL) to the pellet to resuspend. Triton 10% (250 μL) was added to the solution and incubated for 15 min. The solution was filtered using 30 μm Miltenyil filters and added to the sucrose cushion. After centrifugation at 5000 rpm for 10 min, the supernatant was carefully removed, and the pellet was washed twice with Phosphate-Buffered Saline (PBS) 1X and 2% Bovine Serum Albumin (BSA; 3.5 mL, followed by centrifugation at 5,000 rpm for 5 min). Finally, the pellet was resuspended in 50 μL PBS 1X + 2% BSA and filtered through an in-house cotton column to remove debris (Lee and Lin, 2005). Nuclei quality was assessed through DAPI and AO/PI staining at 20x and 40x magnification. Nuclei quantification was performed with a fluorescent cell counter LUNA-FL (Logos Biosystem, Inc., South Korea).

### snRNA-seq library preparation and sequencing

The nuclei suspension was diluted to 1,000 nuclei/μL using 1X PBS + 2% BSA. Single nuclei were partitioned and barcoded using the Chromium Next GEM Single Cell 3’ Reagent Kit V4.0 (10x Genomics). A total of 15 samples (four for control and infected at one dpi; three for control and four for infected at five dpi) were processed randomly in three batches. Libraries were created according to the manufacturer’s protocol. Sequencing was performed on NovaSeq X plus 10B or 25B flow cell, using 150-bp paired-end sequencing, with an average sequencing depth of 40,000 reads per nucleus. Of the 15 samples, 12 (three biological replicates per treatment) were used for the grapevine single-nucleus atlas (**Fig. S1A)**, and eight infected samples (four per one- and five- dpi) were used for *E. necator* (**Fig. S1C**).

### snRNA-seq analysis

For the alignment of snRNA-seq data, a custom reference genome was generated by concatenation of the genomes of *V. vinifera* cv. Cabernet Sauvignon clone FPS 08 v. 2.0 (https://www.grapegenomics.com/pages/VvCabSauv/VvCabSauv08_v2.0/; Cochetel et al., 2025), *E. necator* (Zaccaron et al., 2023), grapevine chloroplast (Arroyo-Garcia et al., 2006) and mitochondria (Goremykin et al., 2009). Reads were mapped using alevin-fry (He et al., 2022) simpleaf function (He and Patro, 2023). The average mapping rate was 92.8 ± 4.25 % (listed per sample in **Table S1**). The count matrix was extracted with loadfry function from the fishpond package in R (Love, 2025). Empty barcodes were filtered out using DropletUtils (Lun et al., 2018). The following analysis was performed with the Seurat package v.5.3.0 (Satija et al., 2015). Low quality nuclei with fewer than 200 and higher than 3,000 reads and with percent mitochondria and chloroplast reads > 10% were discarded. Nuclei with percent *E. necator* reads > 10% were used for the pathogen single-nuclei transcriptomic analysis as described later. Genes expressed in less than three nuclei were discarded. To account for batch effect between samples, we integrated the data using FindIntegrationAnchors to identify the anchors and IntegrateData (with normalization.method = “SCT”). Data was normalized using SCTransform, followed by an initial clustering using RunPCA (with 5,000 variable genes and 30 principal components (PCs)), FindNeighbors, and FindClusters (resolution 0.6). Data was visualized with non-linear dimensionality reduction uniform manifold approximation and projection (UMAP; Healy and McInnes, 2024) using RunUMAP. For cluster annotation, a list of marker genes was made using information from the literature (Baumgart et al., 2025), FindAllMarkers on raw expression data (with logfc.threshold = 0.1, min.pct = .05, only.pos = T, test.use = ‘wilcox’) and FindMarkers between each combination of clusters. Cell type markers were filtered to p_val_adj < 0.05. Cell type marker genes were visualized with the DotPlot function, using parameter scale=T, which scales the average expression values across all genes to have a mean of 0 and variance of 1 within each gene.

### Pseudobulk snRNA-seq analysis

The gene expression of main defense-related gene sets (from Massonnet et al., 2022) was averaged using the function AddModuleScore (Hao et al., 2021) in Seurat and visualized with non-linear dimensionality reduction UMAP. We calculated the 95% confidence intervals (CI) of the module score estimated marginal means per each cluster and treatment. Pseudobulk analysis per cell type was performed using the function AggregateExpression to sum read counts for each cell type and sample (Tenorio Berrío et al., 2025). The pseudobulk differential expression analysis was performed with limma (Hao et al., 2021) to account for zero-inflated data (Lun et al., 2016), to calculate the expression fold change given by the PM infection at each time point of the experiment for each cell type. Gene ontology enrichment analysis on the biological processes was performed using the topGO function in R and the *V. vinifera* PN40024 v5.1 annotation (https://grapedia.org/genomes/). Significant biological processes were defined by *p* value <0.01. Synteny comparison between *V. vinifera* PN40024 (Shi et al., 2023) and Cabernet Sauvignon clone 08 (Cochetel et al., 2025) was performed using MCscan (Wang et al., 2024) from the JCVI v1.0.9 toolkit (Tang et al., 2024).

### Gene co-expression analysis

Gene co-expression networks were calculated and visualized using high-dimensional weighted gene co-expression network analysis with default parameters (hdWGCNA; Morabito et al., 2023). hdWGCNA constructs a gene-gene correlation adjacency matrix to infer co-expression relationships between genes. To account for the complexity and the varied correlation structure of single-nuclei data for different subsets (cell types, cell states, anatomical regions), we collapsed highly similar cells into “metacells” to reduce sparsity while retaining cellular heterogeneity and by allowing for a modular design to perform network analyses in specified cell populations. Metacells were calculated using the function MetacellsByGroups (which applies K-nearest neighbors (KNN) approach to a dimensionality-reduced representation of the dataset), grouping nuclei by samples and cell types, using UMAP dimensionality reduction, k=20 and maximum number of shared cells between metacells of 10. The gene-gene correlations are raised to a power to reduce the amount of noise present in the correlation matrix, thereby retaining the strong connections and removing the weak connections. We determined a proper value for the soft power threshold by picking the lowest soft power threshold that has a Scale Free Topology Model Fit greater than or equal to 0.8 (**Fig. S2A**). We constructed the co-expression network using the default parameters of the ConstructNetwork function. Networks were constructed using the metacell expression matrix from the epidermis, but we computed module eigengenes (first principal component of the gene expression matrix for each module) using the function ModuleEigengene on the entire snRNA-seq dataset, allowing us to interrogate the cell-type specificity of these modules’ expression programs across all cell types (Han et al., 2024; Morabito et al., 2023). We retrieved the harmonized module eigengenes (hME) using the getME function. We computed module eigengene-based connectivity (kME; pairwise correlation between genes and modules) using the ModuleConnectivity function. The top genes with higher kME were considered hub genes and were visualized with the function PlotKMEs (**Fig. S2B**). The gene expression score for the top 25 hub genes per each module was calculated using the UCell method (Andreatta and Carmona, 2021).

### *Erysiphe necator* single-nucleus atlas

Droplets containing *E. necator* reads were processed further. First, to select the droplets with a high likelihood of being fungal nuclei, we calculated quantiles of the proportion of fungal features in each nucleus. The 95^th^ quantile was selected, which included nuclei with ≥ 73.5% of fungal features (). These nuclei were then subjected to additional filtering to remove potential outliers like doublets or artifacts. For this, a locally estimated scatterplot smoothing (LOESS) model (Jacoby, 2000) was fitted to the relationship between the number of fungal reads and the proportion of fungal features per nucleus. Nuclei with residual exceeding a robust Z-score threshold (|Z| > 2.5) were excluded, resulting in a final set of 3,219 nuclei. Data was normalized using SCTransform, followed by an initial clustering using RunPCA (with 2,000 variable genes and 50 PCs), FindNeighbors, and FindClusters (resolution 0.3). Data was visualized with non-linear dimensionality reduction UMAP (Healy and McInnes, 2024) using RunUMAP with 15 dimensions. FindAllMarkers (with logfc.threshold = 1, min.pct = .05, only.pos = F, test.use = ‘wilcox’) was used in combination with manual selection to detect new potential markers for each of the clusters detected. The functional annotation of the reference fungal genes was obtained from Zaccaron et al (2023), and complemented with InterProScan v.5.69-101.0 with default parameters. The full functional annotation of the *E. necator* reference genome used in the analysis can be retrieved in **Table S2.** Additionally, hdWGCNA was applied to the fungus dataset following the method used for the grape presented above. The soft power selected in this dataset was 8.

### Total RNA from bulk leaf tissue

Total RNA isolation was isolated using a Cetyltrimethylammonium Bromide (CTAB)-based protocol as described in Blanco-Ulate et al. (2013) with slight modifications. Isopropanol was used to precipitate both large and small RNAs, and an additional clean-up step was included using the Quick RNA miniprep kit (Zymo Research, Irvine, USA) following the manufacturer’s protocol. Total RNA concentration and purity were assessed using a Qubit fluorometer (Life Technologies, Carlsbad, CA, USA) and a NanoDrop One spectrophotometer (Thermo Scientific, MA, USA), respectively.

### Bulk RNA-seq libraries

RNA-seq libraries were prepared using the Illumina TruSeq RNA sample preparation kit v.2 (Illumina, CA, USA), following Illumina’s Low-throughput protocol. Final libraries were evaluated for quantity and quality with the High Sensitivity chip in a Bioanalyzer 2100 (Agilent Technologies, CA, USA). Libraries were sequenced on the Elements Bio Aviti platform as 80-bp paired-end reads (DNA Technology Core Facility, University of California, Davis, CA, USA).

### Bulk RNA-seq analysis

The Nextflow pipeline nf-core/rnaseq v.3.12.0 (https://github.com/nf-core/rnaseq/tree/3.12.0; 10.5281/zenodo.1400710) was used to process the RNA-seq data. Adaptor and quality trimming were performed with TrimGalore v.0.6.7 (Krueger, 2015), and reads were aligned to the combined transcriptome of Cabernet Sauvignon clone FPS08 and *E. necator* using Salmon v.1.10.1 (Patro et al., 2017). Quantification was conducted with Salmon using the parameters --seqBias and -- posBias. Differential expression analysis was performed with DESeq2 (Love et al., 2014) with the parameter design = ∼ Condition + Timepoint + Condition:Timepoint to get the gene expression fold change given by the infection at each time point of the experiment.

## Data availability

Sequencing data are accessible through NCBI under the BioProject PRJNA1295527.

## Code availability

The analysis pipeline used in this study is available in https://github.com/mariasoleb/Dual_SN_GenExpr_VitisEnecator

## Acknowledgments

We acknowledge the valuable assistance from Alexandra Ramirez Celis for troubleshooting the nuclei isolation protocol, Hong Qiu for the 10x Genomics library preparation, and Bradley Saburo Shibata for the SEM imaging sample preparation.

## Author contributions

M.-S.B., and D.C. designed the project. D.C. supervised the project and secured the funds. M-S.B. carried out all experiments and nuclei extractions. R.F.-B. produced all sequencing libraries. M.-S.B., M.M., M.Z. isolated and maintained the pathogen strain. M.-S.B. performed the data analyses. J.G. analyzed the pathogen gene expression. N.C. supported the bioinformatic analysis. M.-S.B., J.G., and D.C. wrote the manuscript.

## References

Aguilar, G., Pedersen, C., Thordal-Christensen, H., 2016. Identification of eight effector candidate genes involved in early aggressiveness of the barley powdery mildew fungus. Plant Pathol. 65, 953–958.

Aguilar, P.S., Engel, A., Walter, P., 2007. The plasma membrane proteins Prm1 and Fig1 ascertain fidelity of membrane fusion during yeast mating. Mol. Biol. Cell 18, 547–556.

Amrine, K.C., Blanco-Ulate, B., Riaz, S., Pap, D., Jones, L., Figueroa-Balderas, R., Walker, M.A., Cantu, D., 2015. Comparative transcriptomics of Central Asian Vitis vinifera accessions reveals distinct defense strategies against powdery mildew. Hortic. Res. 2.

Andreatta, M., Carmona, S.J., 2021. UCell: Robust and scalable single-cell gene signature scoring. Comput. Struct. Biotechnol. J. 19, 3796–3798.

Arroyo-García, R., Ruiz-García, L., Bolling, L., Ocete, R., López, M., Arnold, C., Ergul, A., Söylemezo Lu, G., Uzun, H., Cabello, F., 2006. Multiple origins of cultivated grapevine (Vitis vinifera L. ssp. sativa) based on chloroplast DNA polymorphisms. Mol. Ecol. 15, 3707–3714.

Arya, G.C., Cohen, H., 2022. The multifaceted roles of fungal cutinases during infection. J. Fungi 8, 199.

Baumgart, L.A., Greenblum, S.I., Morales-Cruz, A., Wang, P., Zhang, Y., Yang, L., Chen, C., Dilworth, D.J., Garretson, A.C., Grosjean, N., 2025. Recruitment, rewiring and deep conservation in flowering plant gene regulation. Nat. Plants 1–14.

Blanco-Ulate, B., Vincenti, E., Powell, A.L., Cantu, D., 2013. Tomato transcriptome and mutant analyses suggest a role for plant stress hormones in the interaction between fruit and Botrytis cinerea. Front. Plant Sci. 4, 142.

Boller, T., Felix, G., 2009. A renaissance of elicitors: perception of microbe-associated molecular patterns and danger signals by pattern-recognition receptors. Annu. Rev. Plant Biol. 60, 379–406.

Both, M., Eckert, S.E., Csukai, M., Müller, E., Dimopoulos, G., Spanu, P.D., 2005. Transcript profiles of Blumeria graminis development during infection reveal a cluster of genes that are potential virulence determinants. Mol. Plant. Microbe Interact. 18, 125–133.

Calonnec, A., Cartolaro, P., Poupot, C., Dubourdieu, D., Darriet, P., 2004. Effects of Uncinula necator on the yield and quality of grapes (Vitis vinifera) and wine. Plant Pathol. 53, 434–445.

Cantu, D., Segovia, V., MacLean, D., Bayles, R., Chen, X., Kamoun, S., Dubcovsky, J., Saunders, D.G., Uauy, C., 2013. Genome analyses of the wheat yellow (stripe) rust pathogen Puccinia striiformis f. sp. tritici reveal polymorphic and haustorial expressed secreted proteins as candidate effectors. BMC Genomics 14, 270.

Chen, D., Hu, H., He, W., Zhang, S., Tang, M., Xiang, S., Liu, C., Cai, X., Hendy, A., Kamran, M., 2022. Endocytic protein Pal1 regulates appressorium formation and is required for full virulence of Magnaporthe oryzae. Mol. Plant Pathol. 23, 133–147.

Chhillar, H., Jo, L., Redkar, A., Kajala, K., Jones, J.D., Ding, P., 2025a. Cell-type-specific execution of effector-triggered immunity. bioRxiv 2025–06.

Chhillar, H., Nguyen, H.H., Yeh, P.-M., Jones, J.D., Ding, P., 2025b. Modular mechanisms of immune priming and growth inhibition mediated by plant effector-triggered immunity. Cell Rep. 44.

Cochetel, N., Vondras, A., Figueroa-Balderas, R., Liou, J., Peluso, P., Cantu, D., 2025. Phased epigenomics and methylation inheritance in a historical Vitis vinifera hybrid. bioRxiv 2025–05.

Delannoy, E., Batardiere, B., Pateyron, S., Soubigou-Taconnat, L., Chiquet, J., Colcombet, J., Lang, J., 2023. Cell specialization and coordination in Arabidopsis leaves upon pathogenic attack revealed by scRNA-seq. Plant Commun. 4.

Douglas, L.M., Wang, H.X., Konopka, J.B., 2013. The MARVEL domain protein Nce102 regulates actin organization and invasive growth of Candida albicans. MBio 4, 10–1128.

Fung, R.W., Gonzalo, M., Fekete, C., Kovacs, L.G., He, Y., Marsh, E., McIntyre, L.M., Schachtman, D.P., Qiu, W., 2008. Powdery mildew induces defense-oriented reprogramming of the transcriptome in a susceptible but not in a resistant grapevine. Plant Physiol. 146, 236–249.

Gadoury, D.M., Cadle-Davidson, L., Wilcox, W.F., Dry, I.B., Seem, R.C., Milgroom, M.G., 2012. Grapevine powdery mildew (Erysiphe necator): a fascinating system for the study of the biology, ecology and epidemiology of an obligate biotroph. Mol. Plant Pathol. 13, 1–16.

Godfrey, D., Böhlenius, H., Pedersen, C., Zhang, Z., Emmersen, J., Thordal-Christensen, H., 2010. Powdery mildew fungal effector candidates share N-terminal Y/F/WxC-motif. BMC Genomics 11, 317.

Goremykin, V.V., Salamini, F., Velasco, R., Viola, R., 2009. Mitochondrial DNA of Vitis vinifera and the issue of rampant horizontal gene transfer. Mol. Biol. Evol. 26, 99–110.

Grell, M.N., Mouritzen, P., Giese, H., 2003. A Blumeria graminis gene family encoding proteins with a C-terminal variable region with homologues in pathogenic fungi. Gene 311, 181–192.

Guo, Y., Yao, S., Yuan, T., Wang, Y., Zhang, D., Tang, W., 2019. The spatiotemporal control of KatG2 catalase-peroxidase contributes to the invasiveness of Fusarium graminearum in host plants. Mol. Plant Pathol. 20, 685–700.

Gururani, M.A., Venkatesh, J., Upadhyaya, C.P., Nookaraju, A., Pandey, S.K., Park, S.W., 2012. Plant disease resistance genes: current status and future directions. Physiol. Mol. Plant Pathol. 78, 51–65.

Han, E., Geng, Z., Qin, Y., Wang, Y., Ma, S., 2024. Single-cell network analysis reveals gene expression programs for Arabidopsis root development and metabolism. Plant Commun. 5.

Hao, Y., Hao, S., Andersen-Nissen, E., Mauck, W.M., Zheng, S., Butler, A., Lee, M.J., Wilk, A.J., Darby, C., Zager, M., 2021. Integrated analysis of multimodal single-cell data. Cell 184, 3573–3587.

He, D., Patro, R., 2023. simpleaf: a simple, flexible, and scalable framework for single-cell data processing using alevin-fry. Bioinformatics 39, btad614.

He, D., Zakeri, M., Sarkar, H., Soneson, C., Srivastava, A., Patro, R., 2022. Alevin-fry unlocks rapid, accurate and memory-frugal quantification of single-cell RNA-seq data. Nat. Methods 19, 316–322.

Healy, J., McInnes, L., 2024. Uniform manifold approximation and projection. Nat. Rev. Methods Primer 4, 82.

Heiman, M.G., Walter, P., 2000. Prm1p, a pheromone-regulated multispanning membrane protein, facilitates plasma membrane fusion during yeast mating. J. Cell Biol. 151, 719–730.

Hu, Z., Chai, J., 2023. Assembly and architecture of NLR resistosomes and inflammasomes. Annu. Rev. Biophys. 52, 207–228.

Huang, S., Jia, A., Ma, S., Sun, Y., Chang, X., Han, Z., Chai, J., 2023. NLR signaling in plants: from resistosomes to second messengers. Trends Biochem. Sci. 48, 776–787.

Jacob, P., Hige, J., Dangl, J.L., 2023. Is localized acquired resistance the mechanism for effector-triggered disease resistance in plants? Nat. Plants 9, 1184–1190.

Jacoby, W.G., 2000. Loess:: a nonparametric, graphical tool for depicting relationships between variables. Elect. Stud. 19, 577–613.

Jiao, C., Sun, X., Yan, X., Xu, X., Yan, Q., Gao, M., Fei, Z., Wang, X., 2021. Grape transcriptome response to powdery mildew infection: comparative transcriptome profiling of Chinese wild grapes provides insights into powdery mildew resistance. Phytopathology® 111, 2041–2051.

Jones, L., Riaz, S., Morales-Cruz, A., Amrine, K.C., McGuire, B., Gubler, W.D., Walker, M.A., Cantu, D., 2014. Adaptive genomic structural variation in the grape powdery mildew pathogen, Erysiphe necator. BMC Genomics 15, 1–18.

Junttila, S., Smolander, J., Elo, L.L., 2022. Benchmarking methods for detecting differential states between conditions from multi-subject single-cell RNA-seq data. Brief. Bioinform. 23, bbac286.

Knepper, C., Savory, E.A., Day, B., 2011. The role of NDR1 in pathogen perception and plant defense signaling. Plant Signal. Behav. 6, 1114–1116.

Krueger, F., 2015. Trim Galore!: A wrapper around Cutadapt and FastQC to consistently apply adapter and quality trimming to FastQ files, with extra functionality for RRBS data. Babraham Inst.

Kunova, A., Pizzatti, C., Saracchi, M., Pasquali, M., Cortesi, P., 2021. Grapevine powdery mildew: fungicides for its management and advances in molecular detection of markers associated with resistance. Microorganisms 9, 1541.

Lee, H.-C., Lin, T.-Y., 2005. Isolation of plant nuclei suitable for flow cytometry from recalcitrant tissue by use of a filtration column. Plant Mol. Biol. Report. 23, 53–58.

Liu, X.-H., Lu, J.-P., Zhang, L., Dong, B., Min, H., Lin, F.-C., 2007. Involvement of a Magnaporthe grisea Serine/Threonine kinase gene, Mg ATG1, in appressorium turgor and pathogenesis. Eukaryot. Cell 6, 997–1005.

Love, M.I., Huber, W., Anders, S., 2014. Moderated estimation of fold change and dispersion for RNA-seq data with DESeq2. Genome Biol. 15, 1–21.

Love, M.M., 2025. Package ‘fishpond.’

Lun, A., Griffiths, J., McCarthy, D., He, D., Patro, R., 2018. DropletUtils: utilities for handling single-cell droplet data. R Package Version 099 14.

Lun, A.T., McCarthy, D.J., Marioni, J.C., 2016. A step-by-step workflow for low-level analysis of single-cell RNA-seq data with Bioconductor. F1000Research 5, 2122.

Massonnet, M., Riaz, S., Pap, D., Figueroa-Balderas, R., Walker, M.A., Cantu, D., 2022. The grape powdery mildew resistance loci Ren2, Ren3, Ren4D, Ren4U, Run1, Run1. 2b, Run2. 1, and Run2. 2 activate different transcriptional responses to Erysiphe necator. Front. Plant Sci. 13, 1096862.

Meng, X., Zhang, S., 2013. MAPK cascades in plant disease resistance signaling. Annu. Rev. Phytopathol. 51, 245–266.

Morabito, S., Reese, F., Rahimzadeh, N., Miyoshi, E., Swarup, V., 2023. hdWGCNA identifies co-expression networks in high-dimensional transcriptomics data. Cell Rep. Methods 3.

Nobori, T., Ecker, J.R., 2023. Yet uninfected? Resolving cell states of plants under pathogen attack. Cell Rep. Methods 3.

Nobori, T., Monell, A., Lee, T.A., Sakata, Y., Shirahama, S., Zhou, J., Nery, J.R., Mine, A., Ecker, J.R., 2025. A rare PRIMER cell state in plant immunity. Nature 638, 197–205.

Novy, V., Carneiro, L.V., Shin, J.H., Larsbrink, J., Olsson, L., 2021. Phylogenetic analysis and in-depth characterization of functionally and structurally diverse CE5 cutinases. J. Biol. Chem. 297, 101302.

Ortiz-Álvarez, J., Becerra-Bracho, A., Méndez-Tenorio, A., Murcia-Garzón, J., Villa-Tanaca, L., Hernández-Rodríguez, C., 2020. Phylogeny, evolution, and potential ecological relationship of cytochrome CYP52 enzymes in Saccharomycetales yeasts. Sci. Rep. 10, 10269.

Paineau, M., Zaccheo, M., Massonnet, M., Cantu, D., 2024. Advances in grape and pathogen genomics toward durable grapevine disease resistance. J. Exp. Bot. erae450.

Patro, R., Duggal, G., Love, M.I., Irizarry, R.A., Kingsford, C., 2017. Salmon provides fast and bias-aware quantification of transcript expression. Nat. Methods 14, 417–419.

Pedersen, C., van Themaat, E.V.L., McGuffin, L.J., Abbott, J.C., Burgis, T.A., Barton, G., Bindschedler, L.V., Lu, X., Maekawa, T., Weßling, R., 2012. Structure and evolution of barley powdery mildew effector candidates. BMC Genomics 13, 694.

Peng, Y.-J., Zhang, H., Wang, G., Feng, M.-G., Ying, S.-H., 2024. MARVEL family proteins contribute to vegetative growth, development, and virulence of the insect fungal pathogen Beauveria bassiana. J. Invertebr. Pathol. 203, 108076.

Pennington, H.G., Jones, R., Kwon, S., Bonciani, G., Thieron, H., Chandler, T., Luong, P., Morgan, S.N., Przydacz, M., Bozkurt, T., 2019. The fungal ribonuclease-like effector protein CSEP0064/BEC1054 represses plant immunity and interferes with degradation of host ribosomal RNA. PLoS Pathog. 15, e1007620.

Qiu, W., Feechan, A., Dry, I., 2015. Current understanding of grapevine defense mechanisms against the biotrophic fungus (Erysiphe necator), the causal agent of powdery mildew disease. Hortic. Res. 2.

Rapicavoli, J.N., Blanco-Ulate, B., Muszyński, A., Figueroa-Balderas, R., Morales-Cruz, A., Azadi, P., Dobruchowska, J.M., Castro, C., Cantu, D., Roper, M.C., 2018. Lipopolysaccharide O-antigen delays plant innate immune recognition of Xylella fastidiosa. Nat. Commun. 9, 390.

Reifschneider, F.J., Boiteux, L.S., 1988. A vacuum-operated settling tower for inoculation of powdery mildew fungi. Phytopathology 78, 1463–1465.

Ritchie, M.E., Phipson, B., Wu, D., Hu, Y., Law, C.W., Shi, W., Smyth, G.K., 2015. limma powers differential expression analyses for RNA-sequencing and microarray studies. Nucleic Acids Res. 43, e47–e47.

Satija, R., Farrell, J.A., Gennert, D., Schier, A.F., Regev, A., 2015. Spatial reconstruction of single-cell gene expression data. Nat. Biotechnol. 33, 495–502.

Sharma, G., Aminedi, R., Saxena, D., Gupta, A., Banerjee, P., Jain, D., Chandran, D., 2019. Effector mining from the Erysiphe pisi haustorial transcriptome identifies novel candidates involved in pea powdery mildew pathogenesis. Mol. Plant Pathol. 20, 1506–1522.

Shi, X., Cao, S., Wang, X., Huang, S., Wang, Y., Liu, Z., Liu, W., Leng, X., Peng, Y., Wang, N., 2023. The complete reference genome for grapevine (Vitis vinifera L.) genetics and breeding. Hortic. Res. 10, uhad061.

Spanu, P.D., 2017. Cereal immunity against powdery mildews targets RNase-Like Proteins associated with Haustoria (RALPH) effectors evolved from a commonancestral gene. New Phytol. 213, 969–971.

Spanu, P.D., Abbott, J.C., Amselem, J., Burgis, T.A., Soanes, D.M., Stüber, K., Loren van Themaat, E.V., Brown, J.K., Butcher, S.A., Gurr, S.J., 2010. Genome expansion and gene loss in powdery mildew fungi reveal tradeoffs in extreme parasitism. Science 330, 1543–1546.

Squair, J.W., Gautier, M., Kathe, C., Anderson, M.A., James, N.D., Hutson, T.H., Hudelle, R., Qaiser, T., Matson, K.J., Barraud, Q., 2021. Confronting false discoveries in single-cell differential expression. Nat. Commun. 12, 5692.

Sree, M.R., Singh, S.K., Prakash, J., Kumar, C., Kumar, A., Mishra, G.P., Sevanthi, A.M., Sreekanth, H., Amala, E., 2024. Powdery mildew pathogen [Erysiphe necator (Schw.) Burrill.] induced physiological and biochemical alterations in leaf tissue of grapevines (Vitis spp.). Physiol. Mol. Plant Pathol. 133, 102386.

Tang, B., Feng, L., Hulin, M.T., Ding, P., Ma, W., 2023. Cell-type-specific responses to fungal infection in plants revealed by single-cell transcriptomics. Cell Host Microbe 31, 1732–1747.

Tang, H., Krishnakumar, V., Zeng, X., Xu, Z., Taranto, A., Lomas, J.S., Zhang, Y., Huang, Y., Wang, Y., Yim, W.C., 2024. JCVI: A versatile toolkit for comparative genomics analysis. Imeta 3, e211.

Tenorio Berrío, R., Verhelst, E., Eekhout, T., Grones, C., De Veylder, L., De Rybel, B., Dubois, M., 2025. Dual and spatially resolved drought responses in the Arabidopsis leaf mesophyll revealed by single-cell transcriptomics. New Phytol. 246, 840–858.

Tsuda, K., Katagiri, F., 2010. Comparing signaling mechanisms engaged in pattern-triggered and effector-triggered immunity. Curr. Opin. Plant Biol. 13, 459–465.

Wang, J., Song, W., Chai, J., 2023. Structure, biochemical function, and signaling mechanism of plant NLRs. Mol. Plant 16, 75–95.

Wang, Jizong, Hu, M., Wang, Jia, Qi, J., Han, Z., Wang, G., Qi, Y., Wang, H.-W., Zhou, J.-M., Chai, J., 2019a. Reconstitution and structure of a plant NLR resistosome conferring immunity. Science 364, eaav5870.

Wang, Jizong, Wang, Jia, Hu, M., Wu, S., Qi, J., Wang, G., Han, Z., Qi, Y., Gao, N., Wang, H.- W., 2019b. Ligand-triggered allosteric ADP release primes a plant NLR complex. Science 364, eaav5868.

Wang, Y., Tang, H., Wang, X., Sun, Y., Joseph, P.V., Paterson, A.H., 2024. Detection of colinear blocks and synteny and evolutionary analyses based on utilization of MCScanX. Nat. Protoc. 19, 2206–2229.

Xue, C., Park, G., Choi, W., Zheng, L., Dean, R.A., Xu, J.-R., 2002. Two novel fungal virulence genes specifically expressed in appressoria of the rice blast fungus. Plant Cell 14, 2107–2119.

Yao, X., Xiong, W., Ye, T., Wu, Y., 2012. Overexpression of the aspartic protease ASPG1 gene confers drought avoidance in Arabidopsis. J. Exp. Bot. 63, 2579–2593.

Zaccaron, A.Z., Neill, T., Corcoran, J., Mahaffee, W.F., Stergiopoulos, I., 2023. A chromosome-scale genome assembly of the grape powdery mildew pathogen Erysiphe necator reveals its genomic architecture and previously unknown features of its biology. Mbio 14, e00645–23.

Zhu, J., Lolle, S., Tang, A., Guel, B., Kvitko, B., Cole, B., Coaker, G., 2023. Single-cell profiling of Arabidopsis leaves to Pseudomonas syringae infection. Cell Rep. 42.

